# Tau load in select brainstem neurons predicts the severity and nature of balance deficits in the absence of cell death

**DOI:** 10.1101/2024.10.14.618073

**Authors:** Yunlu Zhu, Hannah Gelnaw, Paige Leary, Rhoshini Raghuraman, Nitika Kamath, Andy Kraja, Jiahuan Liu, Qing Bai, Shin-ichi Higashijima, Edward A. Burton, David Schoppik

**Affiliations:** Departments of Otolaryngology, Neuroscience & Physiology, and the Neuroscience Institute, New York University Grossman School of Medicine; University of Pittsburgh School of Medicine, Department of Neurology, Pittsburgh, PA; National Institute for Basic Biology, Okazaki, Aichi 444-8787, Japan; Research Center on Life and Living Systems (ExCELLS), Okazaki, Aichi 444-8787, Japan; Pittsburgh VA Healthcare System, Geriatric Research Education and Clinical Center, Pittsburgh, PA

## Abstract

Patients with tauopathies present with profoundly different clinical symptoms^1^, even within the same disorder^2^. A central hypothesis in the field, well-supported by biomarker studies^3,4^ and post-mortem pathology^5–7^, is that clinical heterogeneity reflects differential degeneration of vulnerable neuronal populations responsible for specific neurological functions. Recent work has revealed mechanisms underlying susceptibility of particular cell types^8–10^, but relating tau load to disrupted behavior — es- pecially before cell death — requires a targeted circuit-level approach. Here we studied two distinct balance behaviors in larval zebrafish^11^ expressing a human 0N/4R-tau allele^12^ in select populations of evolutionarily-conserved and well-characterized brainstem vestibular circuits^13,14^. We observed that human tau load predicted the severity of circuit-specific deficits in posture and navigation in the ab- sence of cell death. Targeting expression to either mid- or hindbrain balance neurons recapitulated these particular deficits in posture and navigation. By parametrically linking tau load in specific neu- rons to early behavioral deficits, our work moves beyond cell type to close the gap between pathological and neurological conceptions of tauopathy.

---

We used two well-validated lines to express human 4R-tau in cell-type specific brainstem balance neurons. First, the *Tg(UAS:tau-2a-nls-mCherry)* reporter line^12^ expresses both wild-type human 0N/4R-tau and a nuclear-localized mCherry fluorophore (Figure 1A), allowing unambiguous identification of tau-expressing (tau^+^) cells. Larval zebrafish that express this 4R-tau allele in all neurons show multiple molecular, bio- chemical, pathological, and neurological characteristics of primary 4-repeat tauopathies^12^, including severe neurodegeneration between 2–7 days post-fertilization (dpf) with a median survival of 8–9 days. To inves- tigate how tau accumulation affects behavior before fatal neuronal loss, we instead restricted expression of 4R-tau to specific brainstem balance neurons. The *Is(nefma:Gal4)* driver line^13,15^ targets spinal-projecting midbrain and hindbrain neurons that express neurofilament medium chain, a marker for neurodegeneration^16^.

**Figure 1:**
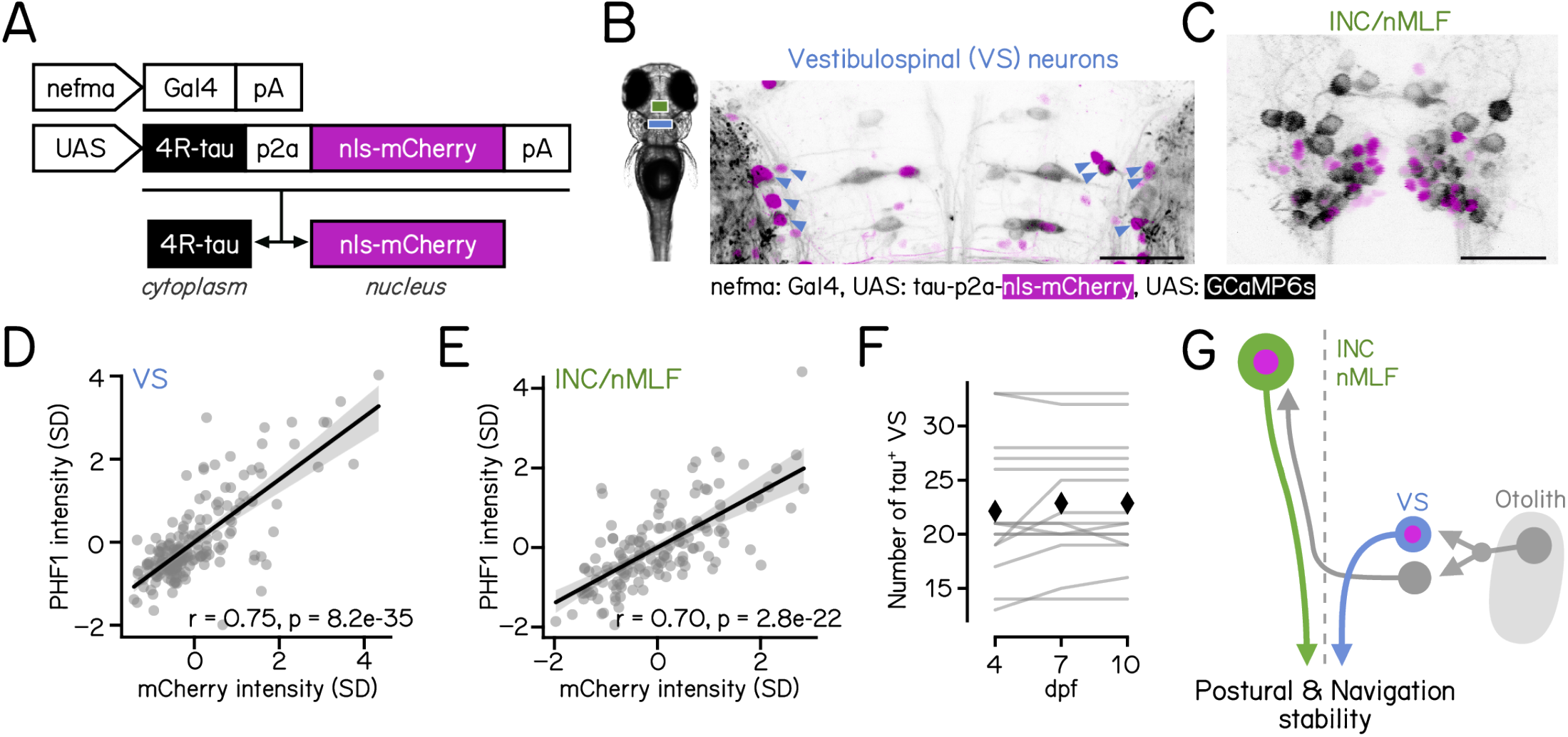
Selective expression of human 0N/4R-tau in brainstem balance neurons without cell death. **(A)** Schematic of the Gal4/UAS alleles used to express cytoplasmic human 4R-tau (black) and nuclear-localized mCherry (nls-mCherry, magenta) in the brainstem using the *nefma* driver line. The p2a peptide allows co-expression of both tau and nls-mCherry ^51^. pA = poly(A) signal. **(B)** Confocal image of the hindbrain (rhombomeres 3–5) from a 7 day post-fertilization (dpf) larvae expressing the *nefma*-driven tau allele (magenta) and cytoplasmic GCaMP6s (greyscale). Blue arrowheads: nuclei from mCherry^+^ vestibulospinal neurons (blue text, VS). Scale bar: 40 µm. **(C)** Confocal image of the midbrain focused on the interstitial nucleus of Cajal / nucleus of the medial longitudinal fasciculus (green text, INC/nMLF, green) from a 7-dpf larvae expressing the *nefma*-driven tau allele (magenta) and GCaMP6s (greyscale). Scale bar: 40 µm. **(D-E)** PHF1 labeling intensity of vestibulospinal (D) and INC/nMLF neurons (E) plotted as a function of their mCherry intensity. Intensity is z-transformed by fish and expressed in units of standard deviation (SD). **(F)** Longitudinal quantification of the number of vestibulospinal (VS) neurons with mCherry^+^ nuclei in each fish. Results from 15 individual fish are shown as gray lines. Diamonds mark the mean across fish. **(G)** Schematic diagram of the direct (blue, VS) and indirect (green, INC/nMLF) balance circuits in the brainstem responsible for postural and navigational behaviors evaluated in this study. The vestibulospinal circuit controls postural stability ^13^ and the INC/nMLF is required for stable navigation ^14^.

We observed tau expression in two evolutionarily-conserved populations of vertebrate brainstem spinal- projecting balance neurons. The first are vestibulospinal neurons in the lateral vestibular nucleus of the hindbrain that are responsible for postural control^13,17^ (Figure 1B). The second are neurons in the midbrain interstitial nucleus of Cajal, also known as the nucleus of the medial longitudinal fasciculus (INC/nMLF, Fig- ure 1C) that are responsible for vertical navigation^14,18^. To confirm expression, we used the antibody PHF1 to label human tau phosphorylated at Ser 396/404^19^. We saw PHF1 signal in the cytoplasm of mCherry^+^ vestibulospinal and INC/nMLF neurons (Figures S1A and S1B). PHF1 intensity and mCherry intensity were strongly correlated (Figures 1D and 1E), establishing that mCherry intensity is an excellent proxy for tau levels. We did not observe PHF1 labeling in mCherry-negative neurons. Importantly, we monitored mCherry intensity in the same cells over time (4–10 dpf) and did not observe neuronal loss (Figure 1F). The absence of cell death allowed us to directly examine the behavioral consequences of 4R-tau expression when restricted to brainstem balance neurons responsible for postural control and navigation (Figure 1G). Notably, as balance is fundamental to locomotion and crucial for survival, larval zebrafish vestibular circuits are online and functional at 4–7 dpf^11,20^ providing an accessible model to resolve the contributions of select groups of neurons to tau-mediated balance deficits.

Posture was unstable in tau^+^ fish. We measured locomotion and posture in the pitch (nose-up/nose-down) axis using an apparatus that recorded sets of 5–7 freely swimming fish from the side^21^. Behavior was recorded in complete darkness when fish were 7–9 dpf. We have previously described how fish maintain posture using swim bouts (Figure 2B) to cancel destabilizing torques^20^. Therefore, we first quantified postural instability as the variability in pitch observed at the end of each swim bout (Figure 2B). Postural instability was always higher in tau^+^ fish compared to control siblings (Figure 2C). Tau-mediated instability was verified on three different genetic backgrounds and with a second 0N/4R-tau allele, *Tg(UAS:tau-2a-EGFP)*; all alleles showed significantly impaired postural stability compared with fish expressing EGFP alone (Figure 2D). Next, we compared the magnitude of instability in tau^+^ fish to changes that followed selective photoablation of vestibulospinal or INC/nMLF neurons. Loss of vestibulospinal neurons led to considerably greater postural instability than any tau line (Figure 2D), consistent with our observation that tau^+^ vestibulospinal neurons remain alive (Figure 1F). We conclude that tau expression in brainstem balance neurons impairs postural stability, and that this impairment is less severe than instability following loss of these neurons.

**Figure 2:**
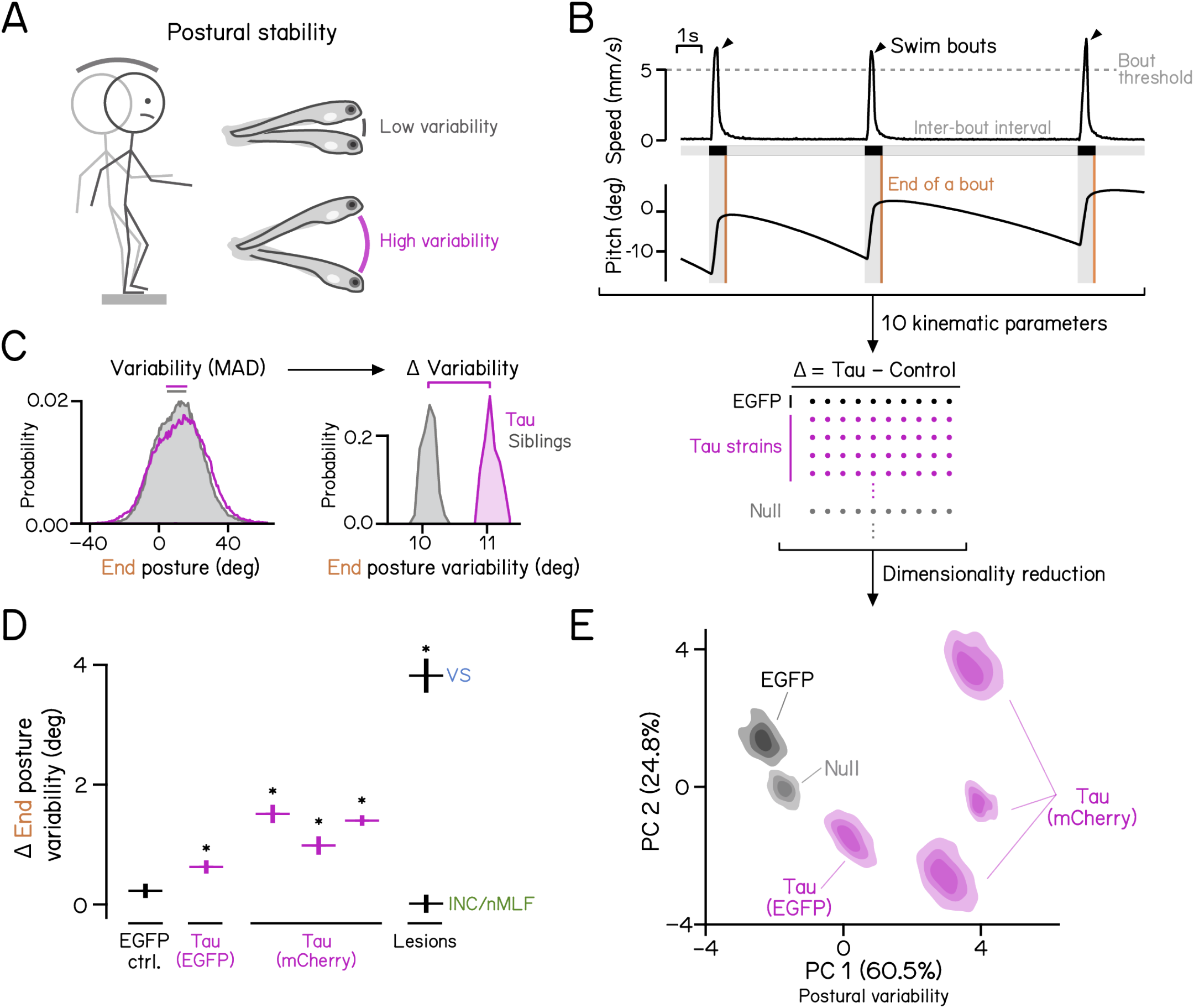
Expression of human 4R-tau in brainstem balance neuronal populations impairs postural stability. **(A)** Schematic illustrating postural stability in the pitch axis (nose-up/nose-down) in humans and fish. Increased postural variability in magenta. **(B)** Behavioral analysis pipeline. Top: A plot of speed (upper) and posture (lower) as a function of time from one larvae. 3 distinct swim bouts (speed *>*5 mm/sec, dashed line) are denoted with arrowheads and gray vertical bars. Gray horizontal bars show the inter-bout interval. Orange lines indicate the end of the swim bout. Bottom: The difference in 10 kinematic parameters (e.g. pitch, speed, and displacement cf. Table 1) between each allele (EGFP control, tau-EGFP, and tau-mCherry on three backgrounds) and sibling controls were used for principal component analysis (PCA). “Null” defined below. Figure S2 illustrates the loadings for each kinematic parameter. **(C)** Left: Distribution of pitch axis posture measured at the end of swim bouts for tau fish (magenta) and sibling controls (gray). Right: Variability of posture was defined as the median absolute difference (MAD) of each distribution. Variance in MAD was estimated by bootstrap resampling. **(D)** Differences (mean ± SD) in pitch axis posture at the end of swim bouts between experimental fish and control siblings. Higher numbers reflect increased variability (degrees). EGFP alone (black), Tau-EGFP (magenta), and Tau-mCherry on three different backgrounds, (magenta) are plotted next to the results of vestibulospinal (VS, blue) and INC/nMLF (green) lesions. A score of 0 represents behavior identical to sibling controls. All conditions were compared to EGFP alone for statistical tests, Tables 2 and 3 for statistics. **(E)** Differences in kinematic parameters between EGFP alone (black), Tau-EGFP (magenta), and Tau-mCherry on three different backgrounds, (magenta) and siblings plotted in PCA space. Contour plots for each allele derived from resampled data. Null refers to a joint distribution drawn from all data to delineate “no effect” in PCA space. Explained variance by each principal component labeled on x and y axes. PC1 loadings encompass measures of postural variability, cf. Figure S2 Refer to Table 1 for parameter definitions, Tables 2 and 3 for parameter values and sample sizes.

**Table 1:** Definitions of behavior parameters. Refer to Figures 2 to 5.

| Parameter/Terminology | Unit | Definition |
| --- | --- | --- |
| Swim bout | – | Period when larvae swim faster than 5 mm/s |
| Inter-bout interval (IBI) | s | Duration between swim bouts |
| *Peak* | – | The time of peak speed within a bout |
| *Initial* | – | 250 ms before the peak speed |
| *End* | – | 200 ms after peak speed |
| Pitch angle | deg | Posture of the fish measured by the pitch (nose-up/down) angle relative to horizontal |
| Dives or climbs | – | Swim bouts where the initial pitch is negative or positive |
| Bout displacement | mm | The Euclidean distance traveled during a bout |
| **10 parameters used for PCA** |  |  |
| Pitch initial variability | deg | MAD of *initial* pitch, representing dispersion of posture just before a bout |
| Pitch peak variability | deg | MAD of *peak* pitch, representing dispersion of posture at peak speed |
| Pitch end variability | deg | MAD of *end* pitch, representing dispersion of posture after a bout |
| IBI pitch variability | deg | MAD of pitch during the IBI |
| Pitch peak dive | deg | Average pitch at peak speed of a dive bout (median) |
| Pitch end dive | deg | Average pitch at the end of a dive bout (median) |
| Pitch peak climb | deg | Average pitch at the end of a climb bout (median) |
| Pitch end climb | deg | Average pitch at the end of a climb bout (median) |
| Swim speed | mm/s | Average peak speed of a bout (median) |
| Displacement | mm | Average displacement of a bout (median) |
| Navigation consistency | – | A measurement of how well swim directions of 4 consecutive bouts are aligned<sup>14</sup>. Refer to Methods for details |

**Table 2:**
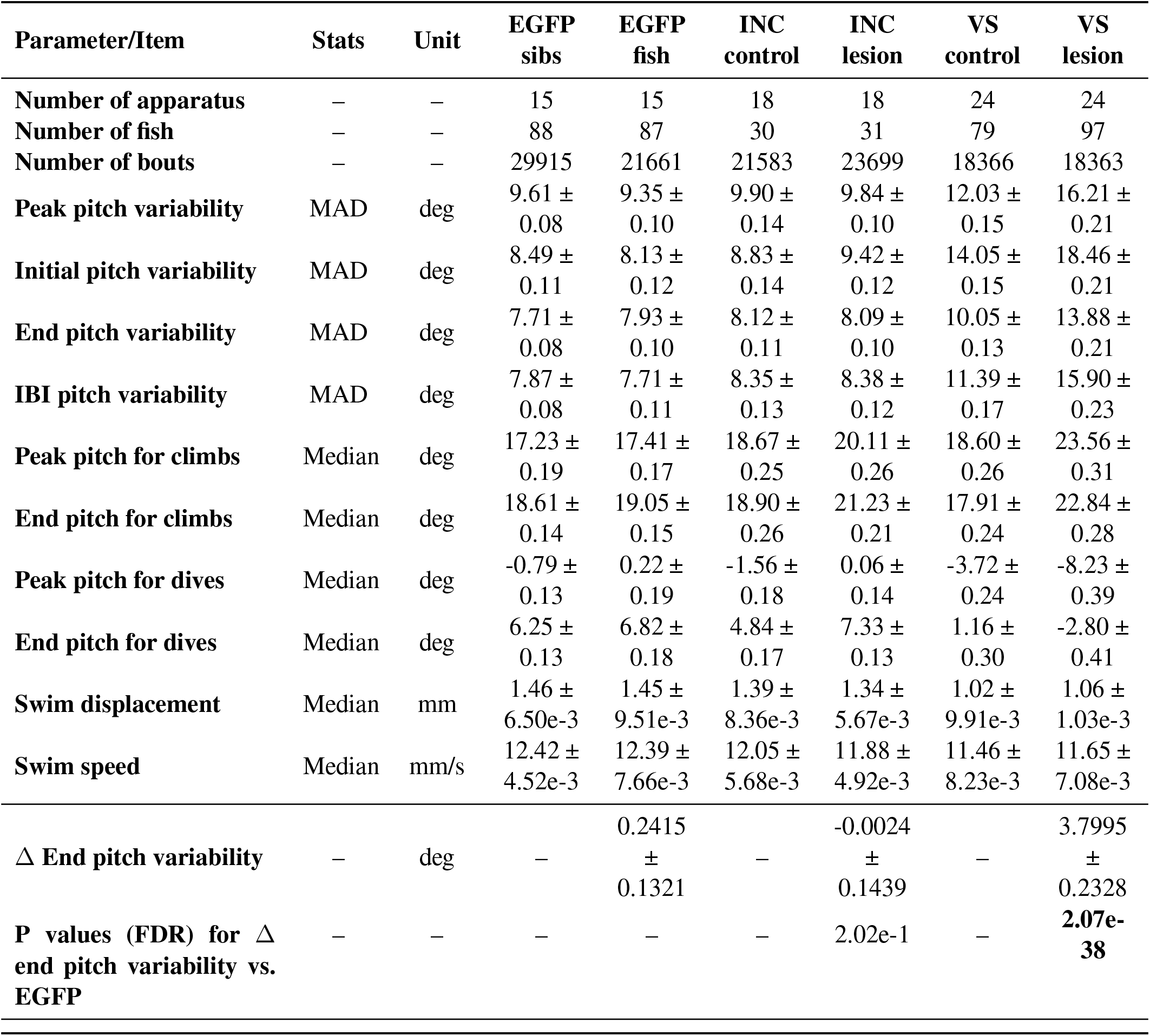
Statistics of swim and postural parameter for EGFP controls and lesion experiments. Refer to Figures 2 and S2. Parameters are calculated as median or median absolute deviation (MAD) across all swim bouts for each condition. The mean ± SD of resampled results are reported.

| Parameter/Item | Stats | Unit | EGFP<br>sibs | EGFP<br>fish | INC<br>control | INC<br>lesion | VS<br>control | VS<br>lesion |
| --- | --- | --- | --- | --- | --- | --- | --- | --- |
| Number of apparatus | – | – | 15 | 15 | 18 | 18 | 24 | 24 |
| Number of fish | – | – | 88 | 87 | 30 | 31 | 79 | 97 |
| Number of bouts | – | – | 29915 | 21661 | 21583 | 23699 | 18366 | 18363 |
| Peak pitch variability | MAD | deg | 9.61 ±<br>0.08 | 9.35 ±<br>0.10 | 9.90 ±<br>0.14 | 9.84 ±<br>0.10 | 12.03 ±<br>0.15 | 16.21 ±<br>0.21 |
| Initial pitch variability | MAD | deg | 8.49 ±<br>0.11 | 8.13 ±<br>0.12 | 8.83 ±<br>0.14 | 9.42 ±<br>0.12 | 14.05 ±<br>0.15 | 18.46 ±<br>0.21 |
| End pitch variability | MAD | deg | 7.71 ±<br>0.08 | 7.93 ±<br>0.10 | 8.12 ±<br>0.11 | 8.09 ±<br>0.10 | 10.05 ±<br>0.13 | 13.88 ±<br>0.21 |
| IBI pitch variability | MAD | deg | 7.87 ±<br>0.08 | 7.71 ±<br>0.11 | 8.35 ±<br>0.13 | 8.38 ±<br>0.12 | 11.39 ±<br>0.17 | 15.90 ±<br>0.23 |
| Peak pitch for climbs | Median | deg | 17.23 ±<br>0.19 | 17.41 ±<br>0.17 | 18.67 ±<br>0.25 | 20.11 ±<br>0.26 | 18.60 ±<br>0.26 | 23.56 ±<br>0.31 |
| End pitch for climbs | Median | deg | 18.61 ±<br>0.14 | 19.05 ±<br>0.15 | 18.90 ±<br>0.26 | 21.23 ±<br>0.21 | 17.91 ±<br>0.24 | 22.84 ±<br>0.28 |
| Peak pitch for dives | Median | deg | -0.79 ±<br>0.13 | 0.22 ±<br>0.19 | -1.56 ±<br>0.18 | 0.06 ±<br>0.14 | -3.72 ±<br>0.24 | -8.23 ±<br>0.39 |
| End pitch for dives | Median | deg | 6.25 ±<br>0.13 | 6.82 ±<br>0.18 | 4.84 ±<br>0.17 | 7.33 ±<br>0.13 | 1.16 ±<br>0.30 | -2.80 ±<br>0.41 |
| Swim displacement | Median | mm | 1.46 ±<br>6.50e-3 | 1.45 ±<br>9.51e-3 | 1.39 ±<br>8.36e-3 | 1.34 ±<br>5.67e-3 | 1.02 ±<br>9.91e-3 | 1.06 ±<br>1.03e-3 |
| Swim speed | Median | mm/s | 12.42 ±<br>4.52e-3 | 12.39 ±<br>7.66e-3 | 12.05 ±<br>5.68e-3 | 11.88 ±<br>4.92e-3 | 11.46 ±<br>8.23e-3 | 11.65 ±<br>7.08e-3 |
| $\Delta$ End pitch variability | – | deg | – | 0.2415<br>±<br>0.1321 | – | -0.0024<br>±<br>0.1439 | – | 3.7995<br>±<br>0.2328 |
| P values (FDR) for $\Delta$ end pitch variability vs. EGFP | – | – | – | – | – | 2.02e-1 | – | **2.07e-38** |

**Table 3:**
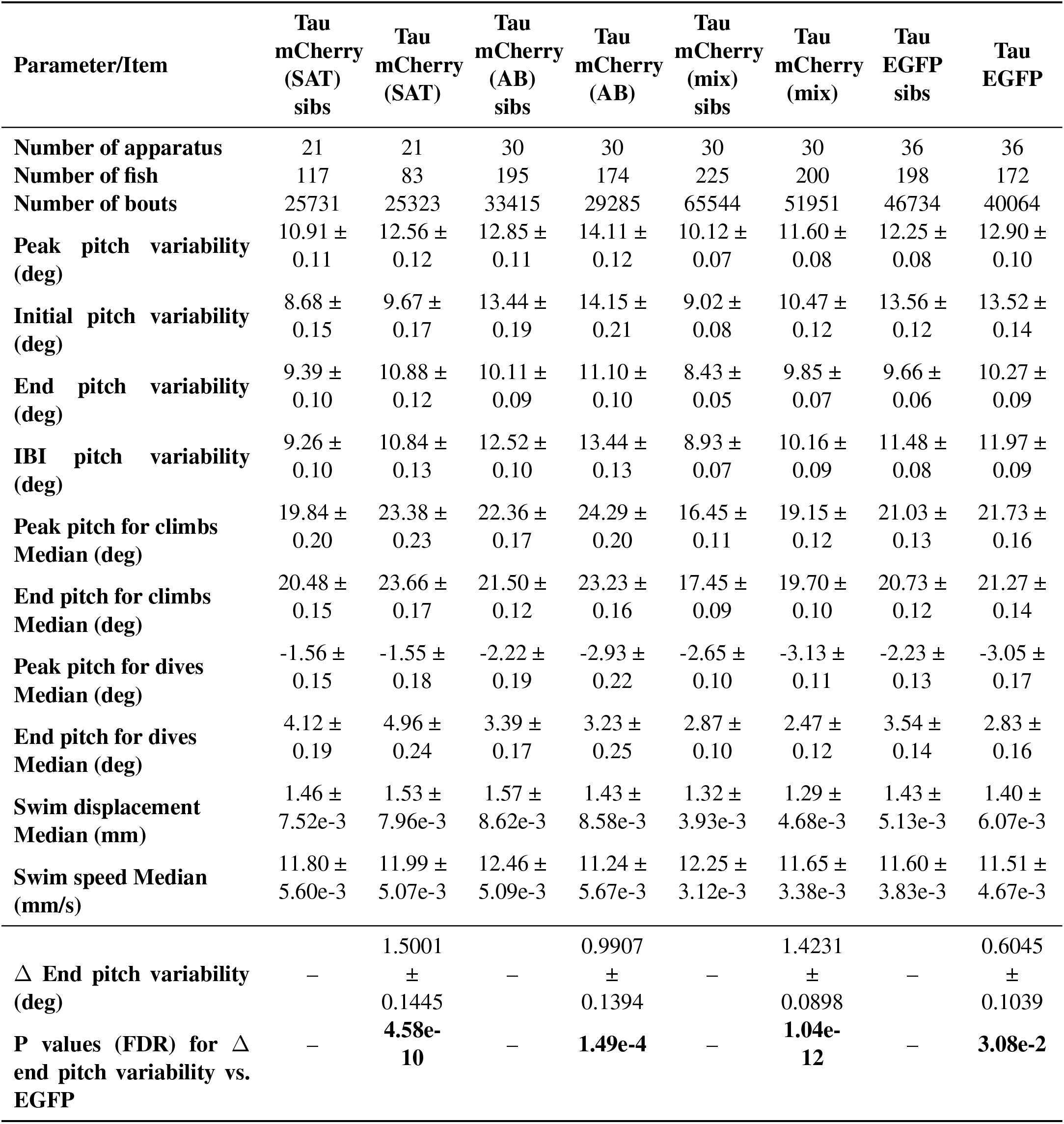
Statistics of swim and postural parameter for Tau fish experiments. Refer toFigures 2 and S2. Parameters are calculated as median or median absolute deviation (MAD) across all swim bouts for each condition. The mean ± SD of resampled results are reported.

An unbiased analysis of changes to postural and locomotor kinematics confirmed that human 4R-tau expression in balance neurons disrupts posture. To move beyond selecting a single variable, we extracted the differences between tau^+^ and control siblings for each of 10 kinematic variables we used previously to characterize and model postural control^20–22^. These differences were then subjected to principal component analysis (PCA, Figure 2B). Consistent with correlations among kinematic variables (Figure S2A), the overwhelming majority (*>*80%) of the variance in our dataset could be explained by two components. The first principal component encompasses postural variability, and the second encompasses locomotor behaviors (Figure S2B). All four tau alleles have more variable posture (PC1) than EGFP-only fish (Figure 2E), but EGFP-only fish sit in the middle of the range of locomotor behaviors (PC2). We conclude that postural stability is selectively impaired in tau^+^ fish.

The variability in expression levels we observed by immunofluorescence (Figures 1D and 1E) suggested a natural experiment that would test whether tau load predicts the severity of postural instability. We measured behavior from single tau^+^ fish, imaged the brainstem by fluorescence microscopy at the end of the experiment, and projected kinematic data on to principal components 1&2 to quantify instability (Figure 3A). Fluorescence intensity was quantified in four regions that encompass key brainstem nodes in postural and navigation control circuits (Figure 3B): vestibulospinal neurons, hindbrain rhombomeres 3–5, the posterior hindbrain, and the INC/nMLF. We then performed multiple linear regression between fluorescent intensity and kinematic data to assess the relationship between tau load and behavioral variability. Fluorescence intensity was moderately correlated across areas but all variance inflation factors^23^ were significantly lower than 5, supporting our regression analysis (Table 4). The intensity in the region with vestibulospinal neurons was the only variable significantly correlated with postural instability (principal component 1, Figure 3C). Next, we simulated the results for a full behavioral experiment (7 fish) where fish had been pre-sorted by vestibulospinal tau load. Consistent with our multiple linear regression, as tau expression increased, so too did postural variability (PC1, Figure 3D) but not locomotion (PC2, Figure 3E). Taken together, we conclude that across individual fish, tau load in vestibulospinal neurons anticipates the severity of postural instability.

**Figure 3:**
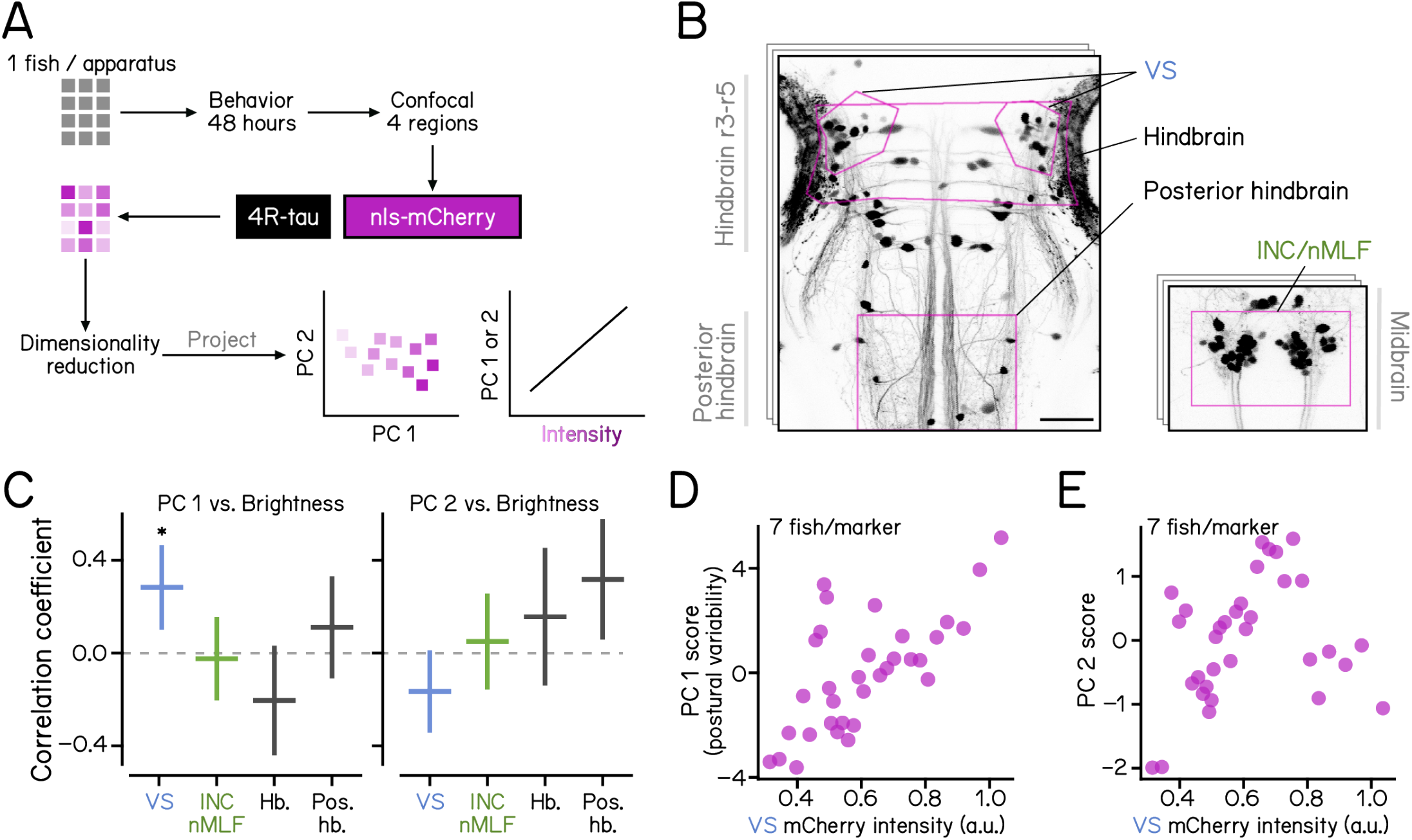
The level of human 4R-tau in vestibulospinal neurons of individual fish predicts postural impairment. **(A)** We recorded behavior from individual fish (1 fish/apparatus, gray square) for 48 hours, took confocal images of mCherry intensity (a proxy for 4R-tau levels) in four regions in the brainstem, then projected single-fish behavioral results into PCA space, and measured correlations between PC1/PC2 values and regional 4R-tau levels. **(B)** A representative example of confocal stacks showing regions of interest used for intensity measurements. Magenta denotes vestibulospinal neurons (VS, blue text), hindbrain, posterior hindbrain, and INC/nMLF (green text, right image). **(C)** Correlation coefficients from multiple linear regression analysis between regional intensity (VS, INC/nMLF, Hindbrain, and Posterior hindbrain) and PC 1 (left) and PC 2 (right) behavioral scores. Dashed lines mark 0. Only vestibulospinal neurons and PC1 were statistically significant. Refer to Table 4 for sample sizes and statistical details. **(D & E)** Each dot represents the behavioral score for PC1 (D) and PC2 (E) for a running average (7 fish wide) arranged in order of increasing brightness in the vestibulospinal region. Single-fish results were calculated using 80264/177231 swim bouts from 39 tau^+^ fish and 126 sibling controls.

**Table 4:**
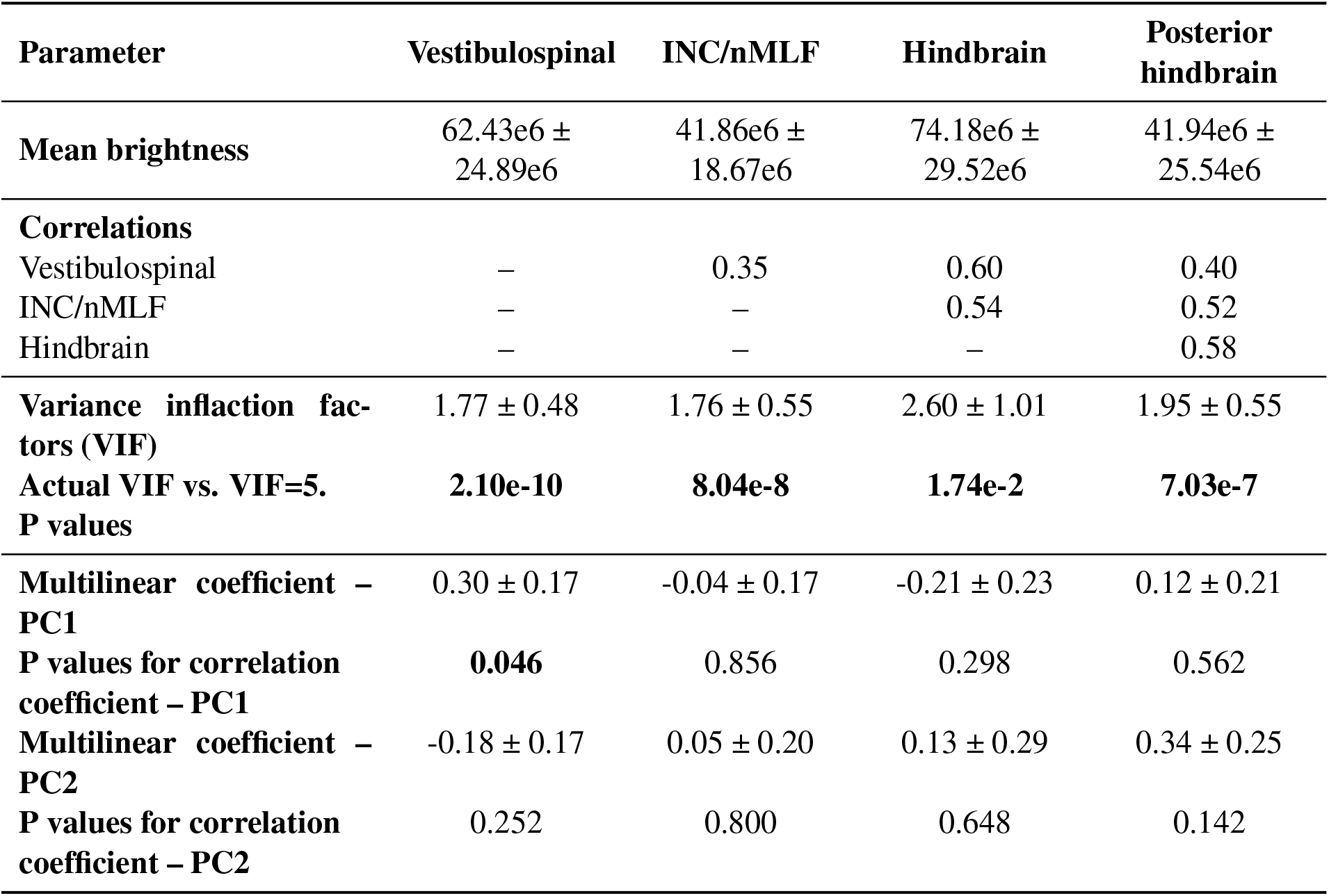
Measurements of mCherry intensity, correlations, and variance inflation factors. Refer to Figure 3. N = 39 fish. The mean ± SD of resampled results are reported.

The ratio of tau load between vestibulospinal neurons and INC/nMLF neurons is positively correlated with disruptions to navigation. Fish navigate in depth by maintaining a consistent heading to arrive at a different elevation (Figures 4A and 4B). Previously, we discovered that INC/nMLF lesions decreased heading consistency, while vestibulospinal lesions stabilized heading^14^ (Figure 4C). Consistency varied among tau^+^ fish: Two out of four allele / genetic backgrounds had significantly more stable navigation than their siblings (Figure 4D). We then computed the ratio of intensity between vestibulospinal neurons and INC/nMLF neurons in single fish (Figure 4E). This intensity ratio was positively correlated with consistency (Figure 4F). Finally, we simulated the results for a full behavioral experiment (7 fish) where fish had been pre-sorted by the intensity ratio. Again, the intensity ratio strongly anticipates the change in heading consistency. We conclude that tau load in vestibulospinal and INC/nMLF neurons together anticipates the severity of navigational instability.

**Figure 4:**
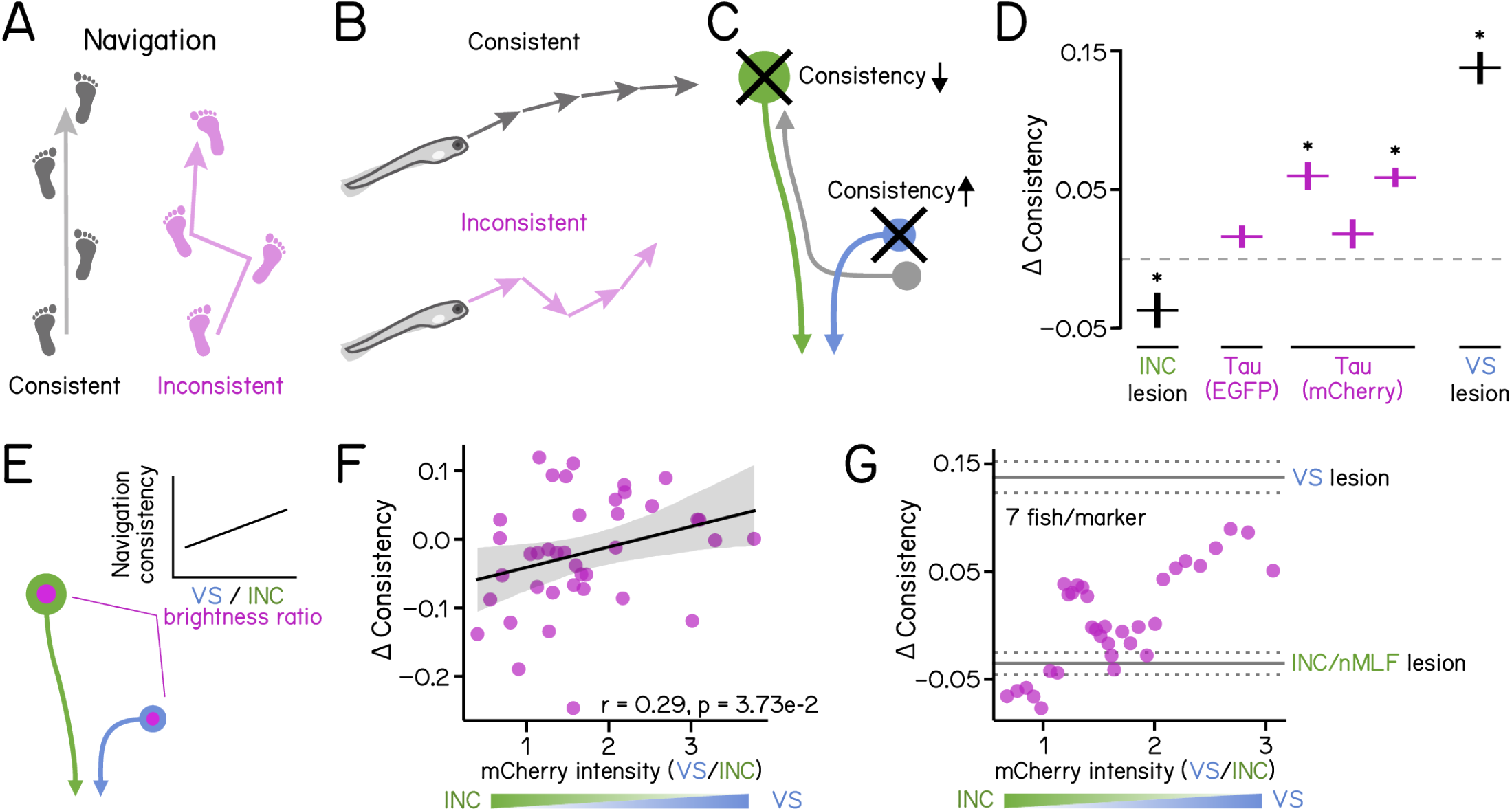
Relative expression of human 4R-tau in vestibulospinal and INC/nMLF neurons predicts the severity of naviga- tional impairment. **(A-B)** Consistent and inconsistent navigation schematized in humans (A) and fish (B). **(C)** Schematic diagram of brainstem bal- ance circuits illustrating differential effects of neuronal loss on navigational consistency: lesion of INC/nMLF (green) neurons reduces consistency while vestibulospinal (blue) ablation increases consistency. Details in ^14^. **(D)** Navigation consistency scores quantified as differences between controls and tau-expressing/lesioned larvae. Results are plotted as mean ± SD for four tau alleles (tau-EGFP, and tau-mCherry on three different backgrounds, magenta) and lesions of vestibulospinal (VS, blue text) or INC/nMLF (green text). The dashed line marks 0. Refer to Table 5 for statistics. **(E)** Schematic diagram illustrating the intensity ratio, de- fined as the relative mCherry intensity in vestibulospinal (blue) vs. INC/nMLF (green), plotted against a measure of navigation consistency **(F)** Navigation consistency scores are quantified as the difference between 39 individual larvae and sibling controls, and plotted against the intensity ratio. Fitted regression line (black) and 95% confidence intervals (grey). **(G)** Averaged navigation consistency scores plotted against the intensity ratio. Each point is the average consistency score of 7 fish, sorted according to brightness ratio. Results of vestibulospinal and INC/nMLF lesions are shown as horizontal lines indicating mean ± SD. Single-fish results were calculated using 39 fish with 436 series of 4 bouts on average for each fish and 16647 series of 4 bouts from 126 sibling controls. Refer to Table 1 for parameter definitions. See also Table 4 for parameter values, fish number, and lesion data size.

**Table 5:** Tau fish navigation consistency phenotype scores and raw parameter values. Refer to Figure 4. The mean ± SD of resampled results are reported.

| Parameter | Tau EGFP | Tau<br>mCherry<br>(SAT) | Tau<br>mCherry<br>(AB) | Tau<br>mCherry<br>(mix) | INC/nMLF<br>lesions | VS lesions |
| --- | --- | --- | --- | --- | --- | --- |
| Navigation<br>consistency (control) | 0.73 ± 0.08 | 0.73 ± 0.08 | 0.77 ± 0.06 | 0.74 ± 0.08 | 0.73 ± 0.07 | 0.62 ± 0.07 |
| Navigation<br>consistency (Tau/lesions) | 0.74 ± 0.08 | 0.79 ± 0.07 | 0.79 ± 0.06 | 0.80 ± 0.05 | 0.69 ± 0.09 | 0.76 ± 0.05 |
| Navigation<br>consistency change | 1.68e-2 ±<br>8.93e-3 | 6.33e-2 ±<br>1.15e-2 | 1.66e-2 ±<br>1.02e-2 | 5.90e-2 ±<br>7.80e-3 | -3.54e-2 ±<br>1.24e-2 | 1.40e-1 ±<br>1.29e-2 |
| P values (FDR) for<br>consistency change vs. 0 | 8.16e-2 | **6.08e-7** | 1.02e-1 | **7.22e-18** | **7.34e-4** | **1.16e-39** |

**Table 6:** Postural stability scores for larvae with restricted Tau expression. Refer to Figure 5. The mean ± SD of resampled results are reported. Refer to Table 8 for pairwise multiple comparisons.

| Parameter | EGFP control | mb Tau | nefma Tau | hb Tau | INC/nMLF lesions | VS lesions |
| --- | --- | --- | --- | --- | --- | --- |
| Number of apparatus (sibs/Tau) | 15/15 | 16/ 16 | 36/36 | 18/ 18 | 18/18 | 24/24 |
| Number of fish (sibs/Tau) | 88/87 | 63/62 | 198/172 | 48/49 | 30/31 | 79/97 |
| Number of bouts (sibs/Tau) | 29915/<br>21661 | 14253/<br>13973 | 46734/<br>40064 | 19110/<br>18067 | 21583/<br>23699 | 18366/<br>18363 |
| Postural variability (PC 1 scores) | -2.05 ±<br>0.18 | -1.64 ±<br>0.28 | -1.10 ±<br>0.15 | 0.96 ± 0.27 | -1.07 ±<br>0.21 | 4.90 ± 0.34 |

**Table 7:** Navigation consistency phenotype scores and raw parameter values for larvae with restricted Tau expression. Refer to Figure 5. The mean ± SD of resampled results are reported. Refer to Table 8 for pairwise multiple comparisons.

| Parameter | INC/nMLF lesions | mb Tau | nefma Tau | hb Tau | VS lesions |
| --- | --- | --- | --- | --- | --- |
| Number of apparatus (sibs/Tau) | 18/18 | 16/ 16 | 36/36 | 18/ 18 | 24/24 |
| Number of fish (sibs/Tau) | 30/31 | 63/62 | 198/172 | 48/49 | 79/97 |
| Number of bouts (sibs/Tau) | 21583/ 23699 | 14253/ 13973 | 46734/ 40064 | 19110/ 18067 | 18366/ 18363 |
| Number of bout series (4 bouts, sibs/Tau) | 4355/4743 | 2425/1975 | 6695/5829 | 3277/3073 | 3554/2920 |
| Navigation consistency (control) | 0.73 ± 0.08 | 0.73 ± 0.09 | 0.72 ± 0.09 | 0.72 ± 0.09 | 0.62 ± 0.07 |
| Navigation consistency (Tau/lesions) | 0.70 ± 0.09 | 0.72 ± 0.10 | 0.74 ± 0.08 | 0.81 ± 0.04 | 0.76 ± 0.05 |
| Change of consistency | -3.693e-2 ± 1.09e-2 | -5.17e-3 ± 1.60e-2 | 1.69e-2 ± 7.45e-3 | 8.32e-2 ± 1.21e-2 | 1.39e-1 ± 1.45e-2 |

**Table 8:** Adjusted p values (FDR) for multiple comparisons. Refer to Figure 5. All p values are explicitly computed from resampled distributions.

| **Figure 5B. PC1 (postural variability)** |  |  |  |  |  |  |
| --- | --- | --- | --- | --- | --- | --- |
|  | **EGFP** | **mb tau** | **Tau (EGFP)** | **hb tau** | **INC/nMLF lesion** | **VS lesion** |
| **EGFP** |  | 2.51e-1 | **3.07e-5** | **2.44e-20** | **3.69e-4** | **1.17e-68** |
| **mb tau** |  |  | 1.27e-1 | **7.99e-12** | 1.17e-1 | **2.23e-50** |
| **Tau (EGFP)** |  |  |  | **1.19e-11** | 9.15e-1 | **5.80e-61** |
| **hb tau** |  |  |  |  | **2.27e-8** | **2.97e-18** |
| **INC/nMLF lesion** |  |  |  |  |  | **3.46e-52** |
| **Figure 5C. Navigation consistency scores** |  |  |  |  |  |  |
|  | **INC/nMLF lesion** | **mb tau** | **Tau (EGFP)** | **hb tau** | **VS lesion** |  |
| **INC/nMLF lesion** |  | 1.17e-1 | **2.79e-4** | **5.41e-12** | **1.01e-20** |  |
| **mb tau** |  |  | 1.17e-1 | **1.71e-5** | **3.21e-14** |  |
| **Tau (EGFP)** |  |  |  | **1.20e-5** | **2.69e-13** |  |
| **hb tau** |  |  |  |  | **7.44e-3** |  |

Selectively restricting tau expression to midbrain or hindbrain populations recapitulates circuit-specific disruptions to posture and navigation. We used Cre/lox recombination together with the *Is(nefma:Gal4)* driver to restrict tau expression to hindbrain rhombomeres 3–5 (vestibulospinal neurons, *Tg(hoxb1a:Cre)*) or to the midbrain (INC/nMLF, *Tg(otx2b:Cre)*, Figure 5A). Vestibulospinal tau^+^ fish showed significantly more postural instability than INC/nMLF tau^+^ fish (Figure 5B). Selective expression of 4R-tau in INC/nMLF decreased navigation consistency, while vestibulospinal tau^+^ fish had increased consistency scores (Figure 5C). We conclude that tau expression in specific populations of brainstem neurons is responsible for disruptions to selected components of balance behavior in the absence of cell death.

**Figure 5:**
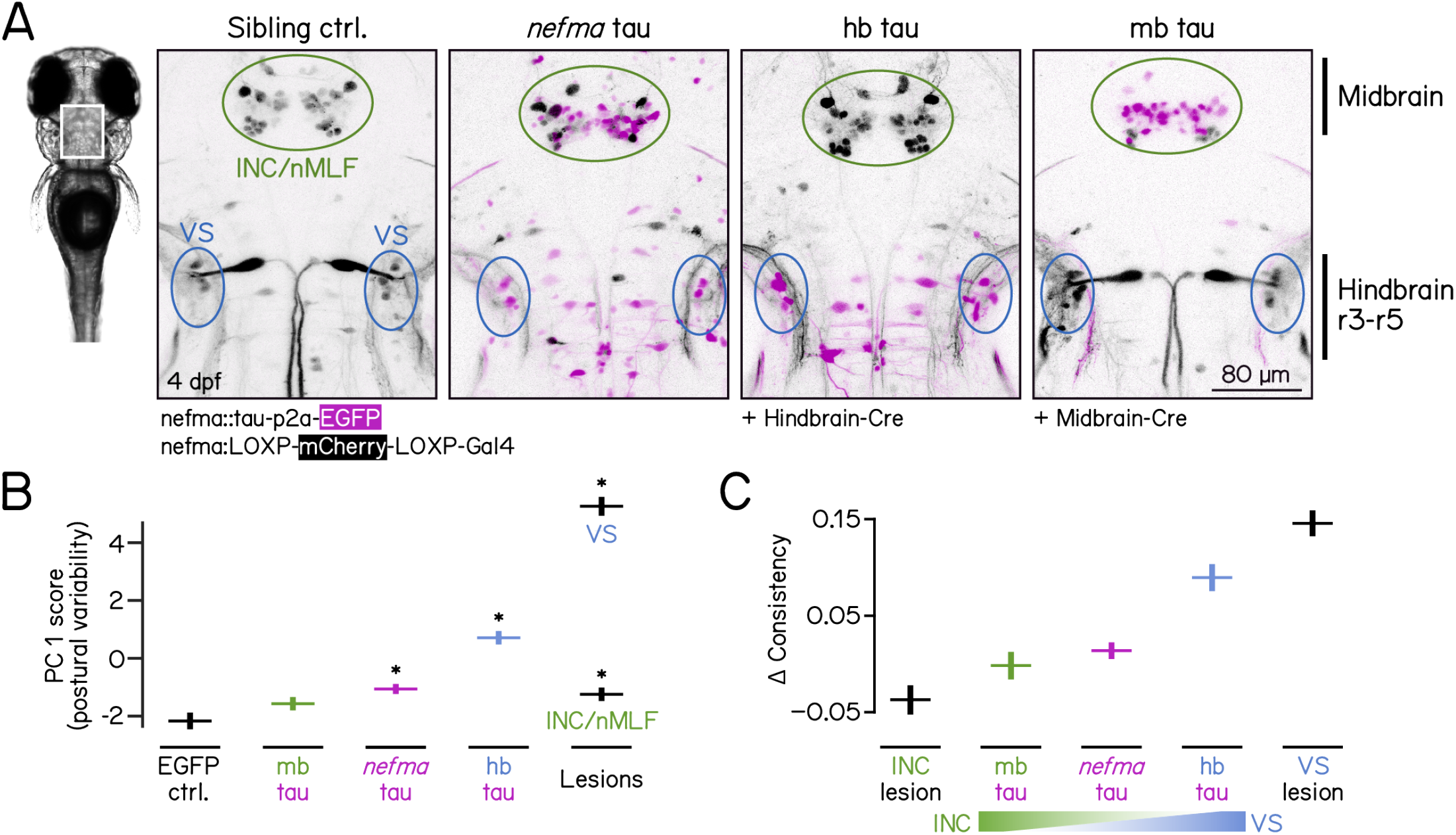
Restricted expression of human 4R-tau selectively recapitulates postural and navigational deficits. **(A)** Confocal images of the brainstem from 4 dpf fish. Sibling controls carrying the unrecombined loxP allele (mCherry, grayscale) show no expression of tau-P2A-EGFP (magenta). *nefma*-Gal4;tau-P2A-EGFP fish show expression of EGFP (magenta) in both hindbrain and midbrain. Cre/lox recombination restricts expression of tau-P2A-EGFP (magenta) to either the hindbrain (hb tau) or midbrain (INC/nMLF tau). Ovals indicate regions of vestibulospinal neurons (VS, blue) and INC/nMLF (green). Scale bar 80 µm **(B)** Postural variability scores (PC1), plotted for EGFP controls (black), *nefma*-Tau expression (magenta), or midbrain/hindbrain restricted expression (green/blue), and lesions of vestibulospinal (VS, blue text) or INC/nMLF (green text). **(C)** Navigation consistency scores plotted for INC/nMLF lesion (black), midbrain tau (mb tau, green), *nefma*-tau (magenta), hindbrain tau (hb tau) and lesions of vestibulospinal neurons (black).

## DISCUSSION

We find that, strikingly, tau expression predicts circuit-specific balance deficits in a load-dependent manner independent of cell death. Load-dependent behavioral disruptions are particularly remarkable given that symptom severity is ordinarily correlated with tau propagation^24^; there was no evidence of tau spread in our model, and the tau^+^ neurons here are roughly one week old^25^. Further, it is notable that we observed a circuit-specific relationship between tau and behavioral disruption. Specifically, regardless of the particular allele or the genetic background, and whether quantification was selective or unbiased, human 0N/4R tau in vestibulospinal neurons disrupts posture, and tau in the INC/nMLF disrupts navigation. Tauopathies are defined with respect to post-mortem pathology; even within a particular disorder, clinical presentation can be heterogeneous^1^. A foundational assumption in the field, supported by correlative post-mortem studies^5,6,26^, is that these heterogeneous symptoms reflect circuit-specific pathophysiology. Our work strongly supports and extends this inference by showing that — even without cell death — tau load parametrically and selectively disrupts behavior in a circuit-dependent manner.

We utilized the 0N/4R-tau isoform, targeted brainstem balance neurons, and modeled balance deficits; all hallmarks of a sporadic primary tauopathy called progressive supranuclear palsy (PSP)^27,28^. Though clinical presentation^29^ and pathological features^6^ are diverse, postural instability^30^ and falls^31,32^ are central to PSP diagnosis. Posturography of PSP patients revealed small but characteristic anteroposterior sway, inconsistent base^33^, and wider step width^34^, with possible parallels to the impaired postural stability (Figure 2A) and navigational consistency (Figures 4A and 4B) we observed in tau^+^ fish. The PSP subtype with predominant postural instability has higher tau load in brainstem regions that encompass the sets of otolith-recipient balance neurons we have studied here^6^. However, processing of otolith-derived information has only been studied in small groups of PSP patients, leaving questions of impairment unresolved^35–37^. Similarly, characterization of postural and locomotor impairments in mouse models of tauopathy (P301S/P301L) has only just begun^38,39^. By linking tau load in brainstem balance neurons to the magnitude of postural and navigational impairment, our work complements and extends both the clinical work and mammalian models of PSP.

Recent advances in molecular profiling^40^ have uncovered putative cellular substrates of neurodegenerative diseases^8,9^ opening new diagnostic and therapeutic avenues^10,41,42^. Cell types are components of the neural circuits that produce behavior. However, circuit-level links between molecular hallmarks of tauopathy and behavioral symptoms have proven elusive. Here we use a cell-type specific driver to restrict human tau expression to key brainstem balance nodes necessary for normal posture and navigation. Rigorous quantification of these balance behaviors revealed a clear link between tau load and dysfunction. By using ancient neural circuits and quantifiable behaviors we bridged the gap between cell type, pathological ground truth and clinical presentation. Similar assays of balance in humans, while considerably more complex, are in-creasingly tractable^43^. Our approach shows how to move beyond cell type to leverage circuit-level insights and behavior to diagnose — and one day treat^44^ — neurodegenerative disease before cell death.

## MATERIALS AND METHODS

### Fish husbandry

All procedures involving larval zebrafish (*Danio rerio*) were approved by the New York University Langone Health Institutional Animal Care & Use Committee (IACUC). Zebrafish embryos and larvae were raised at 28.5°C on a standard 14:10 h light:dark cycle with the lights on from 9 AM to 11 PM. Larvae were raised at a density of 20–50 in 25–40 ml of E3 medium in 10 cm Petri dishes before 5 days post-fertilization (dpf). After 5 dpf, larvae were maintained at densities under 30 larvae per 10 cm petri dish and were fed cultured rotifers (Reed Mariculture) daily.

### Fish lines

*Tg(UAS:tau-p2a-nls-mCherry)* refers to a previously established and validated line^12^, *Tg(UAS:Hsa.MAPT- p2a-nls-mCherry)^pt^*^433^. *Tg(UAS:tau-p2a-EGFP)* refers to a new allele we generated, *Tg(UAS:Hsa.MAPT- p2a-EGFP)^nyc^*^1352^ using the human microtubule associated protein tau mRNA sequence (NCBI reference: NM 016834.5). Its cDNA sequence was fused to p2a-EGFP through a Gly-Ser-Gly link and inserted into a zebrafish Tol2 vector after a 5xUAS promoter (VectorBuilder). The plasmid was injected into *Is(nefma:Gal4)* embryos at the single-cell stage. Injected larvae were screened for EGFP expression in the brainstem at 3 dpf, and the founder was then outcrossed to establish a stable line; F3 fish were used for experiments.

*Is(nefma:Gal4)* refers to *Tg(hsp70l:LOXP-RFP-LOXP-Gal4)^stl^*^601^*^Tg^* ^15^. *Is(nefma:Gal4)*;*Tg(UAS:tau-p2a-nls-mCherry)* zebrafish were crossed to wild types of three different backgrounds — AB, SAT, and a lab strain with mixed AB/TU/WIK background — for behavioral experiments. Photoablations of vestibulospinal neurons and neurons of the INC/nMLF were performed using the *Is(nefma:Gal4)*;*Tg(UAS:EGFP)* on the mitfa^-/-^ background. Lesion datasets were adapted from Zhu et al., 2024^14^.

Because *Tg(hsp70l:LOXP-RFP-LOXP-Gal4)^stl601Tg^* has been constitutively recombined, we used the original allele, *Tg(hsp70l:LOXP-RFP-LOXP-Gal4)^nyc^*^1227^ together with two Cre driver lines to restrict expression to *nefma*^+^ neurons in either the hindbrain and midbrain. Hindbrain expression in rhombomeres 3-5 was achieved with *Tg(hoxb1a-SCP1:BGi-Cre-2A-Cerulean)^y461Tg^* ^45^, abbreviated *Tg(hoxb1a:Cre)*. Midbrain expression was achieved with a new allele, *Tg(otx2b-hs:Cre)^nyc^*^1238^, abbreviated *Tg(otx2b:Cre)*. *Tg(otx2b:Cre)* fish were generated using the CRISPR/Cas9-mediated knock-in method with the hsp70 promoter^46^. The sgRNA sequence for targeting the gene was GGAACCCGGCTAATTGTCTCAGG.

### Measurement of behavior

An extensive description of the materials and methods used to assay posture and locomotion is available in^21^. Briefly, after screening for fluorescence, larvae were transferred to rectangular chambers where they could swim freely while their behavior was monitored and analyzed in real-time. For population behavior assays, each chamber was loaded with either a set of 5–7 tau^+^ larvae or sibling controls. For single fish experiments, either one tau^+^ larvae or 5–7 sibling controls were loaded into each chamber. All behavioral assays were started when fish were 7 days post-fertilization (dpf) between 9 AM and noon, and lasted for approximately 48 hours in complete darkness. After 24 hours of recording (8 dpf), programs were paused for 30 minutes for feeding, during which 1–2 ml of rotifer culture was added to each chamber. Data during the circadian day (9AM–11PM) were used for all analyses.

### Analysis of behavior

All initial kinematic analyses were performed using our previously published pipeline^21^. Briefly, swim bouts are defined as the period when fish translocate faster than 5 mm/s. Inter-bout intervals are the periods between two adjacent swim bouts. Swim bouts were aligned at the time of the peak speed for subsequent analyses.

Pitch is defined as the angle between the horizon and the long axis of the fish’s body; positive values are nose-up. We first aggregated all bouts for a given condition (EGFP, Tau-EGFP, Tau-SAT, Tau-AB, Tau-WT) together. A set of 10 kinematic variables were then extracted from the bout data (Table 1). Variables included median speed, median displacement, and measures of dispersion (median absolute deviation) for bout and inter- bout pitch. As the pectoral fins are engaged when climbing but not diving^22^, bouts were categorized into climbs/dives based on whether the pitch angle before a bout was greater than 0. We then extracted the average (median) pitch at the peak and end of climb and dive bouts.

We next used principal component analysis (PCA) to define the effects of tau expression on behavior. PCA was performed on the difference in each of the 10 kinematic parameters (Table 1) between tau (or EGFP) and control siblings for each condition. Further, to determine the variance in our estimates, we resampled the bout data with replacement 100 times. We generated these resampled datasets for each parameter for each condition and its corresponding controls. We then calculated the differences between control and tau^+^ fish for each parameter in each of the 100 resampled distributions. To allow for comparison across conditions, we z- scored (i.e. subtracted the mean and normalized by the standard deviation) the differences in each parameter. In addition, we generated a “null” dataset by shuffling the data within each of the datasets (EGFP, Tau-EGFP, Tau-SAT, Tau-AB, Tau-WT) and repeating the procedure above. The z-scored values for each parameter for each condition and the null condition were then used for PCA. Data from sets of fish with restricted expression were first analyzed the same way (i.e. differences from control computed, then z-scored). To parameterize the effect of tau expression, data were projected onto the first and second eigenvectors.

Calculation of navigation consistency has been described in detail and validated^14^. Briefly, we extracted bout series that consisted of 4 consecutive bouts. We plotted swim direction, defined by the angle of translocation at the time of the peak speed, of the n^th^ bout in the sequence as a function of the direction of the first bout. The slope of the best-fit line defines the consistency of the n^th^ bout. We averaged consistency values for bouts 2–4 in the series as the navigation consistency score. To estimate the variance in the distribution of swim direction consistency, we resampled the bout series data 100 times with replacement and re-calculated the score.

### Analysis of single-fish behavior

Methods for behavioral recordings of individual fish are the same as for sets of fish. To ensure reliable behavioral measurements, we only used data from chambers that yielded more than 1000 swim bouts. 39/63 tau^+^ fish and 106/126 sibling controls passed the bout number threshold. As above, kinematic parameters were measured. Data from all control fish was pooled to generate a single estimate for each parameter. The differences from the pooled control values were calculated for each tau^+^ fish, then z-scored and projected on to the first two eigenvectors. Projection values for each fish were used as the target variables for multiple linear regression (ordinary least squares model). The predictor values were measurements of fluorescence intensity in different regions, described below. Both the target and predictor variables were z-scored before regression analysis. To estimate the variance in our correlation coefficients, we resampled the data by drawing sets of 39 fish with replacement 1000 times and repeating the procedure above. Finally, we calculated the heading consistency for single fish as described above.

To estimate the behavioral outcome of running a full apparatus (7 fish) with similar intensity measurements, we used a boxcar averaging procedure to combine bout data from those with the closest measurements. Specifically, we sorted fish by their intensity in the vestibulospinal region and aggregated the data from the first 7 fish (1–7), then the next seven (2–8), until (32–39). We then used the combined bout data to repeat the analysis procedure above.

### Quantitative confocal microscopy

All images in the study were taken on a Zeiss LSM 800 equipped with a 20x/1.0 Plan-Apochromat water- immersion objective.

For longitudinal imaging, the same fish were repeatedly imaged at 4, 7, and 10 dpf. For each imaging session, larvae were anesthetized with 0.2 mg/ml Tricane, mounted in 2% low-melting-point agarose, imaged, unmounted, and transferred to 6-well Petri dishes.

For single-fish experiments, larvae were anesthetized at 9 dpf (after measuring behavior) and imaged. Imag- ing settings were held constant across fish. Two image stacks were captured per fish: one at the midbrain covering INC/nMLF and another at the hindbrain that includes reticulospinal neurons and posterior hind- brain. All stacks were 239.5 µm x 319.5 µm, with an interval of 1.5 µm between slices. Regions of interest (ROIs) were hand-drawn for each image. We quantified regional mCherry intensity in Fiji/ImageJ^47^ using the following pipeline: First, to minimize noise, we set subthreshold pixels to NaN by: duplicating the original image, applying a Gaussian blur with sigma = 1 across slices, setting gamma to 0.5 across slices, converting the image to 32-bit, applying a threshold using the “moment preserving” method^48^, setting pixels below the threshold to NaN, and then setting the same pixels in the original image to NaN using image calculation. Next, we quantified regional intensity for each slice in the stack by measuring the median intensity and the number of pixels (area) that were not NaNs for each ROI per slice. The median intensity values were mul- tiplied by the area to generate total intensity per slice. Finally, total intensity values were summed for each ROI across slices to generate the final measurement.

### Whole-mount PHF1 staining and analysis

*Is(nefma:Gal4)*, *Tg(UAS:4R-tau-p2a-nls-mCherry)*, *Tg(UAS:EGFP)* and *Is(nefma:Gal4)*, *Tg(UAS:4R-tau- p2a-nls-mCherry)* larvae were fixed at 7 dpf with 4% PFA in PBS with 1% Triton X-100 for staining. Fixed samples were washed for 5 min using PBS with 1% Triton X-100 (PBSTx) followed by a second 5 min wash using DI water with 1% Triton. Next, fish were permeabilized using acetone, first at room temperature for 5 min, then at -20°C for 10–20 min. Three 5 min PBSTx washes were then performed to clean up acetone. Samples were then blocked and stained with the PHF1 antibody (1:600, a kind gift from Peter Davies) overnight. The following day, samples were washed with PBSTx and stained with a secondary anti- mouse Alexa Fluor 633-labeled antibody (1:600) overnight. On day 3, samples were washed with PBSTx and stored in 50% glycerol in PBS until imaging.

PHF1-stained larvae were mounted in 2% low-melting-point agarose for imaging. PHF1 staining (i.e. Alexa- 633) and mCherry intensity were quantified by drawing an ROI for each cell of interest corresponding to the size of their cell bodies and taking the mean intensity within the ROIs. To account for signal variations across fish caused by the antibody staining procedures, we applied z-standardization to intensity measurements from each fish before concatenating the results.

### Statistics

We used standard bootstrap methods^49^ for statistical analysis of behavioral data. Two-tailed p-values were calculated explicitly using density functions for all comparisons, detailed in Tables 2 to 5 and 8. The Ben- jamini–Hochberg method^50^ was then applied with a false discovery rate of 0.05 to adjust p-values for multiple comparisons. We used multiple linear regression (ordinary least squares) to model the relationship between fluorescent intensity and behavioral phenotype. An ordinary least squares linear regression was used to calcu- late navigation consistency. We used a robust estimator (Theil-Sen) to minimize the effects of outliers when computing navigation consistency scores for individual fish. Finally, Pearson correlation analysis was used to determine relationship between PHF1 intensity and mCherry intensity and to relate changes of consistency scores to intensity ratios.

## Data and Code Availability

All raw data and code are available at the Open Science Framework: doi 10.17605/OSF.IO/7CY5T

## ACKNOWLEDGMENTS

Research was supported by the National Institute of Neurological Disorders and Stroke of the National Institutes of Health under the award numbers R61NS125280 and R33NS125280 (DS and EAB), the National Institute on Deafness and Communication Disorders of the National Institutes of Health under award F31DC020910 (PL), by the Leon Levy Foundation (YZ) and the Rainwater Charitable Foundation (YZ), by the National Science Foundation under Graduate Research Fellowship number DGE2041775 (PL), and by the United States Department of Veterans Affairs under award number BX003168 (EAB). The contents of this article do not represent the views of the United States Government. The authors would like to thank Einar Sigurdsson along with the members of the Schoppik and Nagel labs for their valuable feedback and discussions. The authors dedicate this manuscript to the memory of Phillip R. Certain, Ph.D..

## AUTHOR CONTRIBUTIONS

Conceptualization: YZ, PL, EAB, and DS. Methodology: YZ, QB, HG, and SH. Investigation: YZ, HG, PL, RR, RK, AK, and JL. Formal Analysis: YZ and HG. Visualization: YZ and DS. Reagents: SH, QB, and EAB. Writing: YZ and DS. Editing: YZ and DS. Supervision: DS. Funding Acquisition: YZ, EAB, and DS.

---

**Figure S1:**
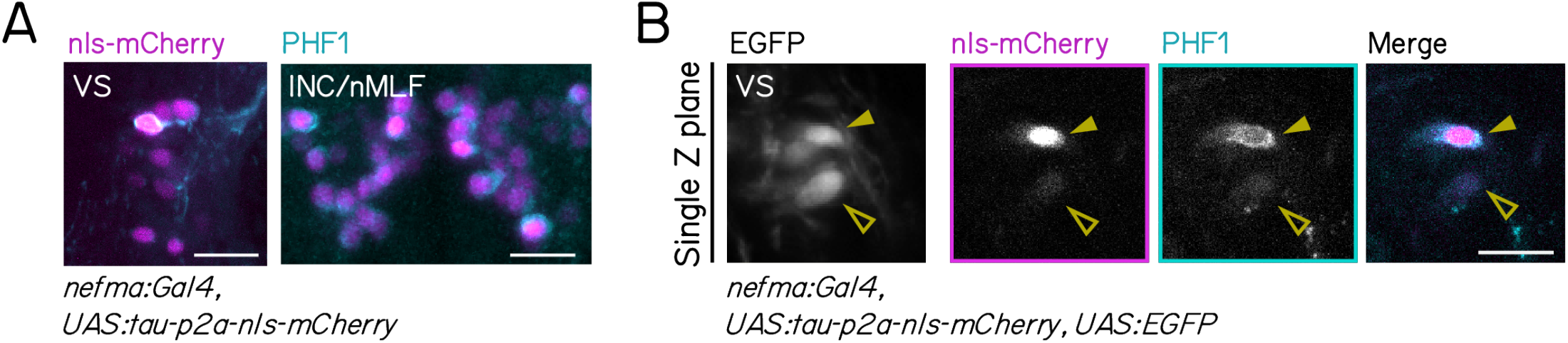
Human 4R-tau is phosphorylated in zebrafish brainstem balance neurons. **(A)** Hindbrain vestibulospinal (VS, left) and midbrain (INC/nMLF, right) of 7 days post-fertilization (dpf) larvae expressing tau- p2A-nls-mCherry (magenta nuclei) stained with PHF1 antibody (cyan cytoplasm) labeling phosphorylated tau. Scale bar: 20 µm. **(B)** Single plane confocal images of vestibulospinal neurons in a 7 dpf tau-p2a-nls-mCherry-expressing larva (magenta) stained with PHF1 antibody (cyan). The vestibulospinal neuron with bright mCherry signal (solid arrowhead) shows stronger PHF1 staining compared to the one with dim mCherry intensity (hollow arrowhead). Scale bar: 20 µm.

**Figure S2:**
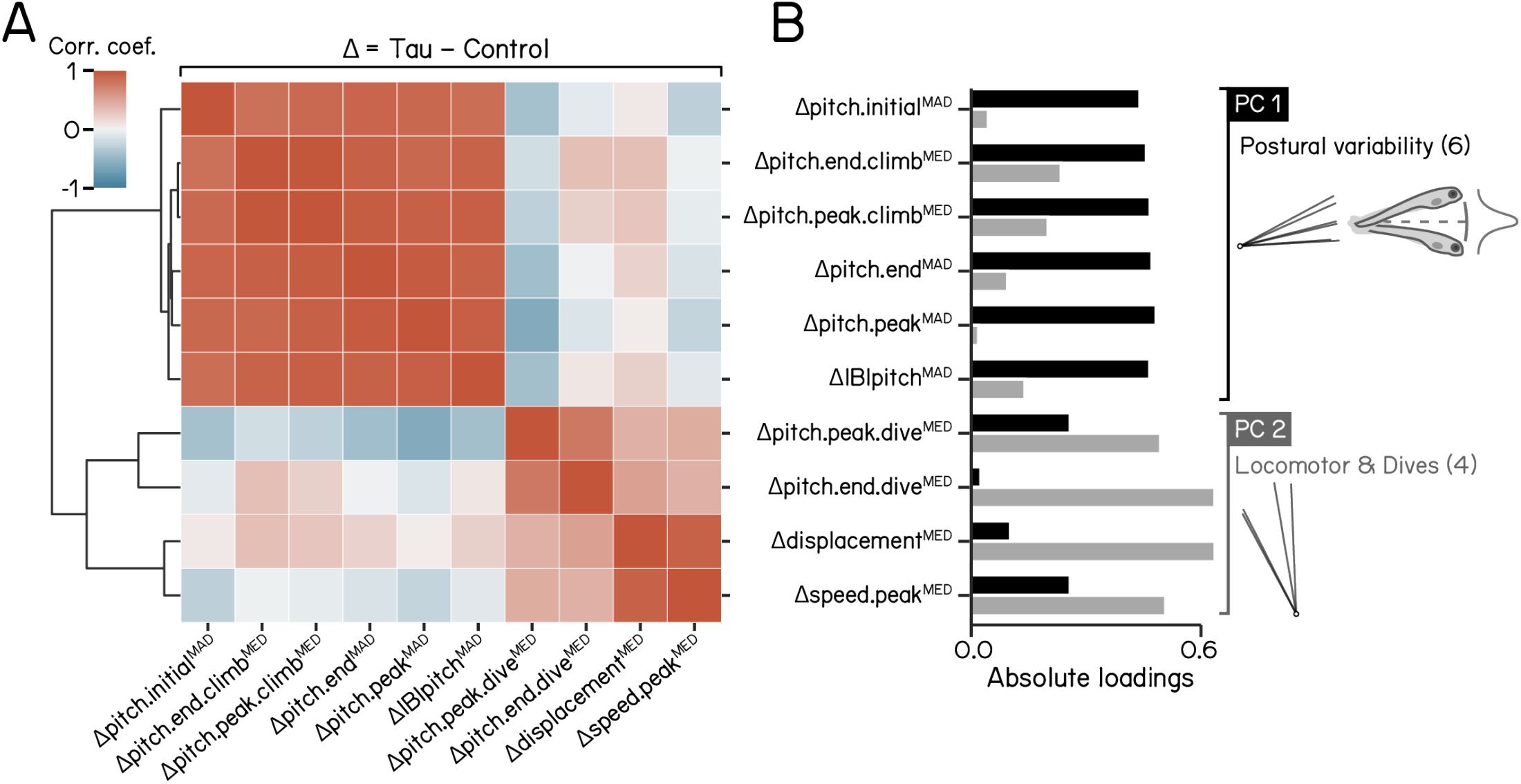
Dimensionality reduction of kinematic differences between tau^+^ fish and control siblings. **(A)** Correlation matrix for changes to kinematic parameters between tau^+^ and sibling controls. A dendrogram is plotted on the left showing clustering of related parameters into two major groups. **(B)** Absolute loadings of differences in kinematic parameters between tau^+^ fish and control siblings for principal components (PC) 1 (black) and 2 (gray). Parameters with strong contributions to each PC were separated and plotted as loading plots on the right, with a schematic illustration of postural variability for PC 1. Refer to Table 1 for parameter definitions, and Tables 2 and 3 for values.

